# Neutralizing antibodies elicited by the Ad26.COV2.S COVID-19 vaccine show reduced activity against 501Y.V2 (B.1.351), despite protection against severe disease by this variant

**DOI:** 10.1101/2021.06.09.447722

**Authors:** Penny L Moore, Thandeka Moyo-Gwete, Tandile Hermanus, Prudence Kgagudi, Frances Ayres, Zanele Makhado, Jerald Sadoff, Mathieu Le Gars, Griet van Roey, Carol Crowther, Nigel Garrett, Linda-Gail Bekker, Lynn Morris, Hanneke Schuitemaker, Glenda Gray

**Affiliations:** National Institute for Communicable Diseases of the National Health Laboratory Service (NHLS), Johannesburg, South Africa; Antibody Immunity Research Unit, School of Pathology, Faculty of Health Sciences, University of the Witwatersrand, Johannesburg, South Africa; Janssen Vaccines and Prevention, Leiden, the Netherlands; Centre for the AIDS Programme of Research in South Africa (CAPRISA), Durban, South Africa; Discipline of Public Health Medicine, School of Nursing and Public Health, University of KwaZulu-Natal, Durban, South Africa; Desmond Tutu HIV Centre, Cape Town, South Africa; South African Medical Research Council, Cape Town, South Africa

## Abstract

The emergence of SARS-CoV-2 variants, such as 501Y.V2, with immune evasion mutations in the spike has resulted in reduced efficacy of several COVID-19 vaccines. However, the efficacy of the Ad26.COV2.S vaccine, when tested in South Africa after the emergence of 501Y.V2, was not adversely impacted. We therefore assessed the binding and neutralization capacity of n=120 South African sera (from Day 29, post-vaccination) from the Janssen phase 3 study, Ensemble. Spike binding assays using both the Wuhan-1 D614G and 501Y.V2 Spikes showed high levels of cross-reactivity. In contrast, in a subset of 27 sera, we observed significantly reduced neutralization of 501Y.V2 compared to Wuhan-1 D614G, with 22/27 (82%) of sera showing no detectable neutralization of 501Y.V2 at Day 29. These data suggest that even low levels of neutralizing antibodies may contribute to protection from moderate/severe disease. In addition, Fc effector function and T cells may play an important role in protection by this vaccine against 501Y.V2.

## Introduction, Results and Conclusions

The SARS-CoV-2 pandemic has resulted in Emergency Use Authorizations (EUA) of several COVID-19 vaccines. These include the Janssen/Johnson and Johnson single-dose Ad26.COV2.S vaccine, an adenovirus 26 vectored vaccine carrying a stabilized SARS-CoV-2 Wuhan-1 spike ^[1]^. Preclinical studies of this vaccine showed protection in macaques, and phase 1/2 clinical trials demonstrated that Ad26.COV2.S induced binding and neutralizing antibodies as well as cellular responses after a single dose ^[2]^. A phase 3 clinical trial of Ad26.COV2.S conducted in 8 countries on three continents demonstrated an average of 66% efficacy against moderate disease and 85% protection against severe disease by Day 28 postvaccination ^[3]^

The emergence of SARS-CoV-2 variants bearing immune evasion mutations has unfortunately compromised the efficacy of several COVID-19 vaccines. The 501Y.V2 (B.1.351 or Beta) variant, discovered first in South Africa and now detected in many regions, contains multiple antibody escape mutations in the Spike protein, and shows significantly reduced sensitivity to sera from convalescent donors and Pfizer, Moderna and AstraZeneca vaccinees ^[4–7]^. Furthermore, the AstraZeneca vaccine showed negligible protection against mild/moderate COVID-19 disease caused by 501Y.V2 in South Africa. This was consistent with pseudovirus neutralization assays showing 84% neutralization resistance of 501Y.V2 to vaccinee sera ^[7]^.

The Janssen phase 3 study, Ensemble, included more than 6,000 participants in South Africa during the period in which 501Y.V2 became the dominant variant ^[3]^. Although there was 95% prevalence of this variant in COVID-19 cases in the South African sub-study trial, this did not adversely impact efficacy, with 64% and 82% vaccine efficacy at Day 28 post vaccination for moderate and severe COVID-19, respectively. We therefore assessed whether the clinical efficacy of Ad26.COV2.S COVID-19 against 501Y.V2 was associated with preserved neutralization capacity against this variant.

We tested 118 blinded serum samples obtained from South Africans enrolled in the phase 3 clinical trial, including 88 vaccinee and 30 placebo participants (3:1 ratio) at Day 1 (prevaccination) and Day 29 (28 days post-vaccination) (**Table S1**). All samples were assayed for binding antibodies by ELISA against the receptor binding domain (RBD), and the Wuhan-1 D614G Spike as described previously ^[5]^. Of the 118 samples (88 of which were vaccinee sera), 86 bound Spike at Day 29, though only 35 (29%) bound RBD. Binding to the two antigens was highly correlated (r=0,8184, p<0.001) (**Fig. 1A**). We also assayed binding to the 501Y.V2 Spike, and although binding was reduced by 1.8-fold, we observed a strong correlation in titers to both proteins (r=0,9683, p<0.001), indicating that the Ad26.COV2.S vaccine triggered cross-reactive binding antibodies, likely largely directed to epitopes outside the RBD (**Fig. 1B**).

**Figure 1:**
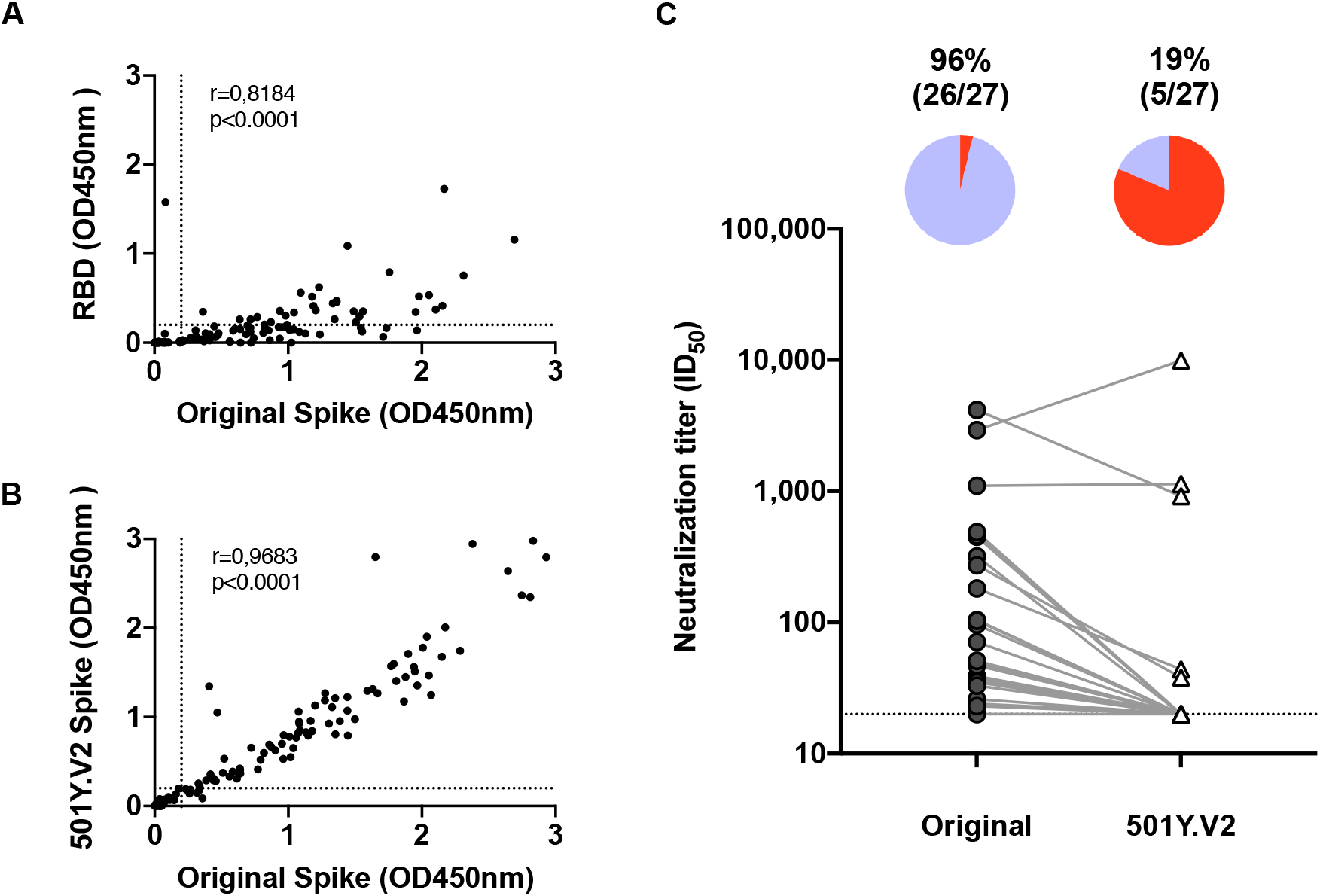
The 501Y.V2 variant is largely resistant to Day 29 neutralizing antibodies elicited by Ad26.COV2.S. (A) Blinded serum samples from 120 participants were screened for binding to receptor binding domain (RBD) and Spike protein from the original variant, with titers showing a strong correlation. (B) A strong correlation was observed in binding titers to the full Spike protein from the original variant and 501Y.V2. (C) A subset of 27 samples from Day 29, positive for binding antibodies to both the original Spike and RBD, were compared for their neutralization of the original variant and 501Y.V2. In the pie charts, purple indicates the proportion of samples with neutralization activity and red the proportion of samples with no detectable neutralization activity. The threshold of detection for the neutralization assay is ID_50_>20. All experiments were performed in duplicate and the average value shown.

We selected all 27 participants that were seronegative at Day 1, but reactive to both the original spike and RBD at Day 29, likely the highest neutralizing responders to Ad26.COV2.S. We used a pseudovirus assay to measure Day 29 neutralization of the original variant and 501Y.V2. Of the 27 sera, 26 (96%) neutralized the Wuhan-1 D614G variant, but only 5 (19%) neutralized 501Y.V2 (**Fig. 1B**), with a 3-fold reduction in geometric mean titer from 152 to 48.

Given the observed vaccine efficacy in South Africa with Ad26.COV2.S, this suggests that even very low levels of neutralizing antibodies at Day 29 may contribute to protection from moderate/severe disease. Moreover, vaccine elicited cross-reactive binding antibodies may harness Fc effector functions against 501Y.V2, and these, along with T cells likely play an important role in protection by this vaccine against 501Y.V2 ^[8,9]^.

## Materials and Methods

### Cohort

A blinded subset of 118 serum samples were obtained from South Africans enrolled in the phase 3 clinical trial of the Ad26.COV2.S COVID-19 vaccine, at a 3:1 ratio of vaccinees to placebo, at Day 1 (pre-vaccination) and Day 29 (28 days post-vaccination). The demographics of the subjects, by vaccine/placebo status after unblinding are summarised in **Table S1**.

### Expression and purification of SARS-CoV-2 antigens

SARS-CoV-2 Spike and RBD (+ subdomain 1) proteins were expressed in Human Embryonic Kidney (HEK) 293F suspension cells by transfecting the cells with SARS CoV-2 plasmid DNA. After incubating for six days at 37 °C, 70% humidity and 10% CO_2_, proteins were first purified using a nickel resin followed by size-exclusion chromatography. Relevant fractions were collected and frozen at −80 °C until use.

### Enzyme-linked Immunosorbent Assay (ELISA)

Two ug/ml of Spike or RBD proteins were coated onto 96-well, high-binding plates and incubated overnight at 4 °C. The plates were incubated in blocking buffer consisting of 5% skimmed milk powder, 0.05% Tween 20, 1x PBS. Serum samples were diluted to 1:100 starting dilution in blocking buffer and 100 ul added to the plates for 75-90 minutes. Secondary antibody was diluted to 1:3000 in blocking buffer and added to the plates for 60 minutes followed by TMB substrate (Thermo Fisher Scientific). Upon stopping the reaction with 1 M H_2_SO_4_, absorbance was measured at a 450nm wavelength. Monoclonal antibodies CR3022 and BD23 were used as positive controls. Palivizumab was used as a negative control.

### SARS-CoV-2 pseudovirus based neutralization assay

SARS-CoV-2 pseudotyped lentiviruses were prepared by co-transfecting the HEK 293T cell line with either the SARS-CoV-2 614G spike (D614G) or SARS-CoV-2 501Y.V2 spike (L18F, D80A, D215G, K417N, E484K, N501Y, D614G, A701V, 242-244 del) plasmids in conjunction with a firefly luciferase encoding lentivirus backbone plasmid and a murine leukemia virus backbone plasmid. The parental plasmids were kindly provided by Drs Elise Landais and Devin Sok (IAVI). For the neutralization assay, heat-inactivated serum samples from vaccinees were incubated with the SARS-CoV-2 pseudotyped virus for 1 hour at 37°C, 5% CO_2_. Subsequently, 1×104 HEK293T cells engineered to over-express ACE-2, kindly provided by Dr Michael Farzan (Scripps Research Institute), were added and incubated at 37°C, 5% CO2_2_ for 72 hours upon which the luminescence of the luciferase gene was measured. Monoclonal antibodies CB6, CA1, 4A8, CC12.1 CR3022 and palivizumab were used as controls.

## Acknowledgements

We acknowledge funding from the South African Medical Research Council (Reference numbers 96825, SHIPNCD 76756 and DST/CON 0250/2012). P.L.M. is supported by the South African Research Chairs Initiative of the Department of Science and Innovation and the National Research Foundation (Grant No 9834). We acknowledge Daniel Stieh for his support in assay discussions and data interpretation. We thank Drs Nicole Doria-Rose, David Montefiori, Elise Landais and Michael Farzan for reagents and assistance in establishing the SARS-CoV-2 pseudotyped neutralization assay and enabling equivalency and proficiency testing. We thank Drs Devin Sok, Elise Landais, Dennis Burton, Nicole Doria-Rose and Peter Kwong for SARS-CoV-2 directed mAbs. We are grateful to Jason McLellan for the WT Hexapro Spike construct and Florian Krammer for the RBD construct.

---

**Table S1:** Summary of Demographics and Baseline Characteristics of Subjects in the South African Subset of the Ad26.COV2.S COVID-19 vaccine trial (Study VAC31518COV3001)

|  | Ad26.COV2.S | Placebo | All subjects |
| --- | --- | --- | --- |
| **Total number [N]** | 88 | 30 | 118 |
| **Age [years]** |  |  |  |
| Mean (SD) | 53.0 (14.80) | 53.7 (15.79) | 53.2 (14.99) |
| Median | 59.5 | 59.5 | 59.5 |
| Range | (23; 77) | (22; 79) | (22; 79) |
| IQ range | (41.0; 64.0) | (41.0; 65.0) | (41.0; 64.0) |
| **Age** |  |  |  |
| [18-39] | 18 (20.5%) | 7 (23.3%) | 25 (21.2%) |
| [40-59] | 26 (29.5%) | 8 (26.7%) | 34 (28.8%) |
| [60-69] | 33 (37.5%) | 11 (36.7%) | 44 (37.3%) |
| [70-79] | 11 (12.5%) | 4 (13.3%) | 15 (12.7%) |
| [>=80] | 0 | 0 | 0 |
| **Sex** |  |  |  |
| Female | 28 (31.8%) | 8 (26.7%) | 36 (30.5%) |
| Male | 60 (68.2%) | 22 (73.3%) | 82 (69.5%) |
| **Race** |  |  |  |
| Asian | 1 (1.1%) | 0 | 1 (0.8%) |
| Black or African American | 70 (79.5%) | 27 (90.0%) | 97 (82.2%) |
| White | 11 (12.5%) | 1 (3.3%) | 12 (10.2%) |
| Multiple | 2 (2.3%) | 1 (3.3%) | 3 (2.5%) |
| Unknown | 2 (2.3%) | 0 | 2 (1.7%) |
| Not reported | 2 (2.3%) | 1 (3.3%) | 3 (2.5%) |
| **SARS-CoV-2 seropositivity status at baseline** |  |  |  |
| Positive | 0 | 0 | 0 |
| Negative | 88 (100.0%) | 30 (100.0%) | 118 (100.0%) |
| Missing | 0 | 0 | 0 |
| **Presence of baseline comorbidity** |  |  |  |
| One or more | 16 (18.2%) | 4 (13.3%) | 20 (16.9%) |
| None | 72 (81.8%) | 26 (86.7%) | 98 (83.1%) |
| **HIV infection** |  |  |  |
| Positive | 2 (2.3%) | 1 (3.3%) | 3 (2.5%) |
| Negative | 14 (15.9%) | 3 (10.0%) | 17 (14.4%) |
| Missing | 72 (81.8%) | 26 (86.7%) | 98 (83.1%) |

